# Identification of Kinase Inhibitors that Regulate Nuclear Receptor Nurr1 (NR4A2) Cellular Activity

**DOI:** 10.1101/420976

**Authors:** Kenya L. Williams, Carrow I. Wells, John T. Moore

**Affiliations:** Shaw University, Division of Science and Technology, Raleigh, NC 27709; SGC-UNC, Eshelman School of Pharmacy, University of North Carolina, Chapel Hill, NC 27599

**Keywords:** NR4A, Nurr1, Nuclear receptor, Kinase inhibitor

## Abstract

The ability to regulate the activity NR4A subfamily of nuclear receptors would be potentially useful in the treatment of multiple diseases, but to date, few regulators have been reported. This is likely due to the fact that the NR4A subfamily does not have a typical unoccupied nuclear receptor ligand binding pocket, but rather a pocket that is occupied with hydrophobic side chains of adjacent amino acids. It follows that traditional nuclear receptor assays that seek to identify ligands that bind within the ligand binding pocket would not be successful. We have thus focused on an alternate assay to identify NR4A regulators based on the fact that regulation of NR4A, at least partially, results from phosphorylation/dephosphorylation of the amino terminal region of the protein. We developed a medium throughput cellular assay using a fusion of the amino terminus of Nurr1 (NR4A2) with luciferase reporter and used the assay to screen a large and diverse protein kinase inhibitor set (PKIS). We identified multiple kinase inhibitor compounds from PKIS that significantly increased or decreased the cellular activity of Nurr1. These molecules serve as starting points to discover selective tools for regulation of Nurr1/Nurr1pathway.

## Introduction

The NR4A proteins (NR4A1/Nur77, NR4A2/Nurr1, and NR4A3/NOR1) are members of the steroid/thyroid superfamily of nuclear receptors (Kurakula *et al*. 2014). These receptors have been identified as potential targets for treatment of multiple diseases (Pearen and Muscat, 2010). This receptor family mediates adaptive and innate immune cell function (e.g., NOR1 is involved in negative selection of thymocytes) (Hamers *et al*., 2013 and Hanna *et al*., 2011), metabolism (e.g., NOR1 augments glucose transport in skeletal muscle cells) (Liu *et al.*, 2013), brain function (e.g., Nurr1 regulates differentiation and maintenance of midbrain dopaminergic neurons) (Sakurada *et al*., 1999) and cancer (e.g., nur77 and NOR1 are tumor suppressors in leukemia) (Beard *et al.*, 2015). The ability to modulate the activity of this receptor family would have significant potential application across multiple human disease areas.

The NR4A subfamily shares a common structural arrangement with other NR family members, possessing an amino terminal region (A/B domain) with a ligand-independent transactivation domain AF1, a DNA-binding domain (DBD), and a C-terminal ligand binding domain (LBD) with an associated AF2 activation domain (Wang *et al*., 2003). Unlike other conventional ligand-regulated NRs, however, the NR4A members lack a hormone binding pocket, with most of the central cavity of their LBD occupied by bulky side chains of three hydrophobic phenylalanine residues (Flaig *et al*., 2005; Davis *et al*., 1993). Thus, these receptors are more likely activated indirectly, through a cell signaling cascade (Berrabah *et al*., 2011) rather than direct binding by a hormone. Finding molecules that regulate NR4A family members is thus likely to require an approach different from the traditional screening the molecules that bind to the ligand binding domain.

There is particular interest in activating Nurr1 for the treatment of Parkinson’s disease (Decressac *et al*., 2013). Nurr1 is expressed in selective brain regions early in development and into adulthood. Specifically, it has been shown to be involved in the establishment and maintenance of the dopaminergic phenotype within central nervous system neuronal subpopulations including the nigrostriatal dopamine system (Chu *et al*., 2006). Clinical and experimental data indicate that disrupted Nurr1 function contributes to induction of midbrain dopaminergic neuron dysfunction, which is seen in early stages of Parkinson’s disease. Nurr1 and its transcriptional targets are down-regulated in midbrain dopaminergic neurons that express high levels of the disease-associated protein α-synuclein. The likely involvement of Nurr1 in the development and progression of Parkinson’s disease makes this protein a potentially interesting target for therapeutic intervention (for review, see Decressae *et al*., 2013). Multiple groups have reported screens for small molecule regulators of Nurr1, but to date, no clinical candidates for this receptor have been progressed.

One high throughput screen identified a single compound, 6-mercaptopurine (6-MP), that acts as an NR4A activator (Ordentlich *et al*., 2003). 6-MP increased the activity of both Nurr1 and NOR1 (6-MP effect on nur77 was difficult to assess due to the high constitutive activity of this receptor). 6-MP is a semisynthetic thiopurine, structurally related to endogenous purine bases adenine, guanine and hypoxanthine. 6-MP, and other thiopurine drugs, were developed in the 1960s for the treatment of leukemia and a number of autoimmune conditions (Elion, 1989; Hitchings and Elion, 2003; Zimmerman *et al*., 1974; Sahasranaman *et al*., 2008). 6-MP depletes cellular purine pools (Nelson *et al*., 1975). Interestingly, studies in CV-1 cells have shown that 6-MP activation of Nurr1 can be blocked by replenishing endogenous purines, adenine, adenosine, guanosine, hypoxanthine, or inosine (Ordentlich *et al*., 2003), consistent with the hypothesis that alteration of cellular purine pools is one of the key events in regulation of Nurr1 by 6-MP.

Deletion studies of Nurr1 showed that 6-MP activates this receptor via the amino-terminal domain, not via a C-terminal Ligand-Binding Domain (Ordentlich *et al*., 2003; Wansa *et al*., 2003). A minimal region in the amino-terminal A/B domain between amino acids 44-151 was essential for activation, a region that contains a ligand-independent transcriptional activation domain (AF1) (Davis *et al*., 1993, Wansa *et al*., 2003). Additional studies on the related NOR1 showed that coactivator recruitment at the amino terminus, specifically TRAP220, was induced by 6-MP and that TRAP220 potentiates transactivation and interacts with the receptor in an AF-1-dependent and cell-specific manner (Wansa *et al*., 2005).

Sequence motifs in the AF1 region that are potential phosphorylation sites are conserved between Nurr-1 and NOR1 (Ordentlich *et al*., 2003). A mutational analysis has demonstrated that NOR1 is a substrate of DNA-dependent protein kinase (DNA-PK) and is phosphorylated in the N-terminal domain in vascular smooth muscle cells (Medunjanin *et al*., 2015). Phosphorylation resulted in post-transcriptional stabilization of the protein through prevention of receptor ubiquitination. In another study, Nurr1 was shown to be phosphorylated by ERK2 and increased phosphorylation resulted in increased tyrosine hydroxylase gene expression in SHSY cells (Zhang *et al*., 2007).

As a step toward characterization of the signaling pathway linking 6-MP to NR4A2 activation, we carried out a broad screen of kinase signaling pathways using the Published Kinase Inhibitor Set (PKIS) (Drewry *et al*., 2014; Elkins *et al*., 2015), which contains 367 small-molecule ATP-competitive kinase inhibitors. To screen the set, we developed a cellular Nurr1 assay using the amino terminal portion of the receptor. The kinase inhibitors discovered in this screen can be used to dissect the cellular pathways involved in regulation of NR4A activity and potentially elucidate new mechanisms to treat NR4A-relevant diseases.

## Materials and Methods

### NR4A2 Transient Transfection assay

Transient transfections were carried out in CV-1 cells (ATCC CCL70) using the NR4A2-AB-Gal4DBD expression plasmid (Li *et al*., 2012, gift from Dr. Stephen Safe, Texas A&M) and the UAS-tk-luciferase reporter. Nurr1-AB-Gal4DBD expresses the Nurr1 A/B domain (amino acids 1-259) fused to the Gal-4 DNA-binding domain, a construct shown to contain a region sufficient for maximal activation by 6-MP (Li *et al*., 2012). Transfection conditions were optimized using the positive control compound (6-MP, 100 μM) versus DMSO. In the final medium throughput assay, transfections were carried out in 96-well format (Costar 96-well solid white plates) with CV-1 cells at approximately 70% confluence. A 12:1 ratio of Fugene-6 (ul) to DNA (ug) was delivered to the cells on Day 1, followed by removal of the transfection mixture and addition of compounds on Day 2. Compound treatment was carried out for 18h under standard cell culture conditions. Effects of compounds on NR4A2-AB-Gal4DBD activity was determined via end-point luciferase measurement using a Luciferase Assay System kit (Promega, Inc). The response to 6-MP (3-4-fold maximal induction, approximate EC_50_ = 80 μM) was similar to that seen in previous studies (Ordentlich *et al.*, 2003). In the medium throughput assay, test compounds were added in a volume of 5ul (<2% DMSO, final concentration) and incubated for 18h under standard cell culture conditions. All transfections were carried out in the presence of 100uM 6-MP except for the negative control wells.

### Cellular ATP assay

The Cell Titer Glo Assay (Promega, Inc) was used to measure cellular ATP levels in a set of parallel assays in the compound screen, as described in manufacturer’s instructions.

#### PKIS screen

PKIS, a set of 367 small molecule kinase inhibitors, was obtained fom the SGC-UNC. PKIS compound structures and associated kinase selectivity data (screening panels represent approximately 200 kinases) are non-proprietary and publically accessible via https://pharmacy.unc.edu/research/sgc-unc/. PKIS was screened in the NR4A2-AB-Gal4DBD transient transfection assay (+/− 100 μM 6-MP, 384-well) at singe shot (10 μM) assays, n = 4, with the positive controls (100 μM 6-MP), and the negative controls (DMSO and a vector-alone control to identify compounds that inhibit end-point luciferase assay in absence of pCMV-NR4A2-Gal4).

## Results

The complete PKIS compound library was screened using our optimized Nurr1 medium throughput assay. “Hits” in the assay were defined as compounds that changed NR4A2 activity > 3-fold the standard deviation difference from the mean, where the mean was defined by replicates of pCMV-NR4A2-Gal4DBD + 6-MP control. Cellular ATP levels were measured in parallel-treated plates. In the absence of any other treatment, the positive control 6-MP significantly reduced cellular ATP levels (30-40%) after the 18-hour incubation, as anticipated from previous reports (Ordentlich *et al*., 2003). The screen was run twice in independent tests and only compounds that were identified as hits in both of the runs are reported. Data was normalized to the 6-MP control (100%) for both Nurr1 activity (luciferase assay) and cellular ATP levels (Cell Titer Glo assay).

The screen identified hits that were fell into the following 3 functional categories: A) Nurr1 activators, B) Nurr1 inhibitors, and C) Nurr1 inhibitors that additionally inhibited the cellular ATP decrease caused by 6-MP.

### Nurr1 Activators

17 compounds activated Nurr1 activity without affecting the cellular ATP decrease caused by 6-MP (Figure 1 **and** Table 1). These compounds fell into 11 chemotypes (Table 1). Assessment of kinase activity data on the compounds showed overlapping kinase targets between chemotypes. For example, KIT was inhibited by three of the chemotypes and GSK3A and GSK3B was inhibited by two chemotypes. Also, interestingly two of the compounds did not identify any kinases with greater than 50% Inhibition, highlighting the fact that current kinase selectivity panels only represent a subset of the human kinome.

**Figure 1.**
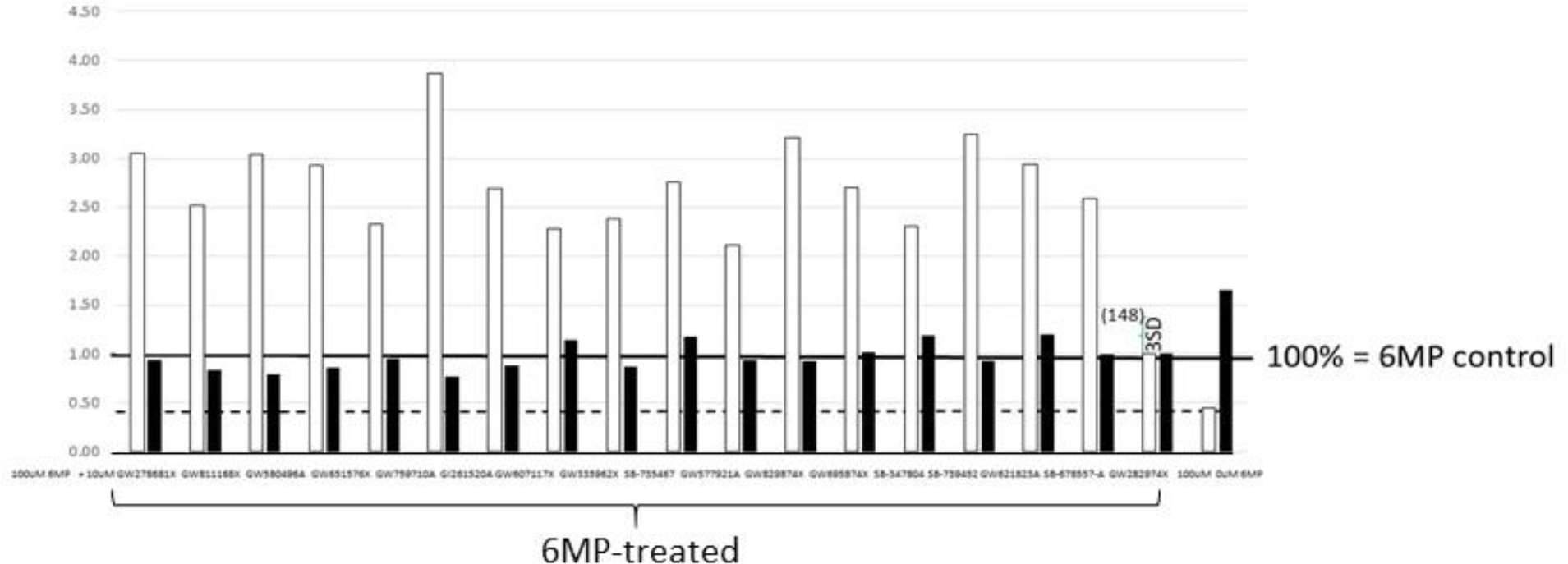
Seventeen compounds activated Nurr1 activity without significantly affecting the cellular ATP decrease caused by 6-MP. Controls are 100uM 6MP and OuM 6MP treatments. Data are normalized to the 100uM 6MP treatment (1.0 = 100%). Three (3) Standard deviations from the 6MP control value illustrated by bracket. White squares represent Nurr1 activityc and Black squares represent cellular ATP levels.

**Table 1.**
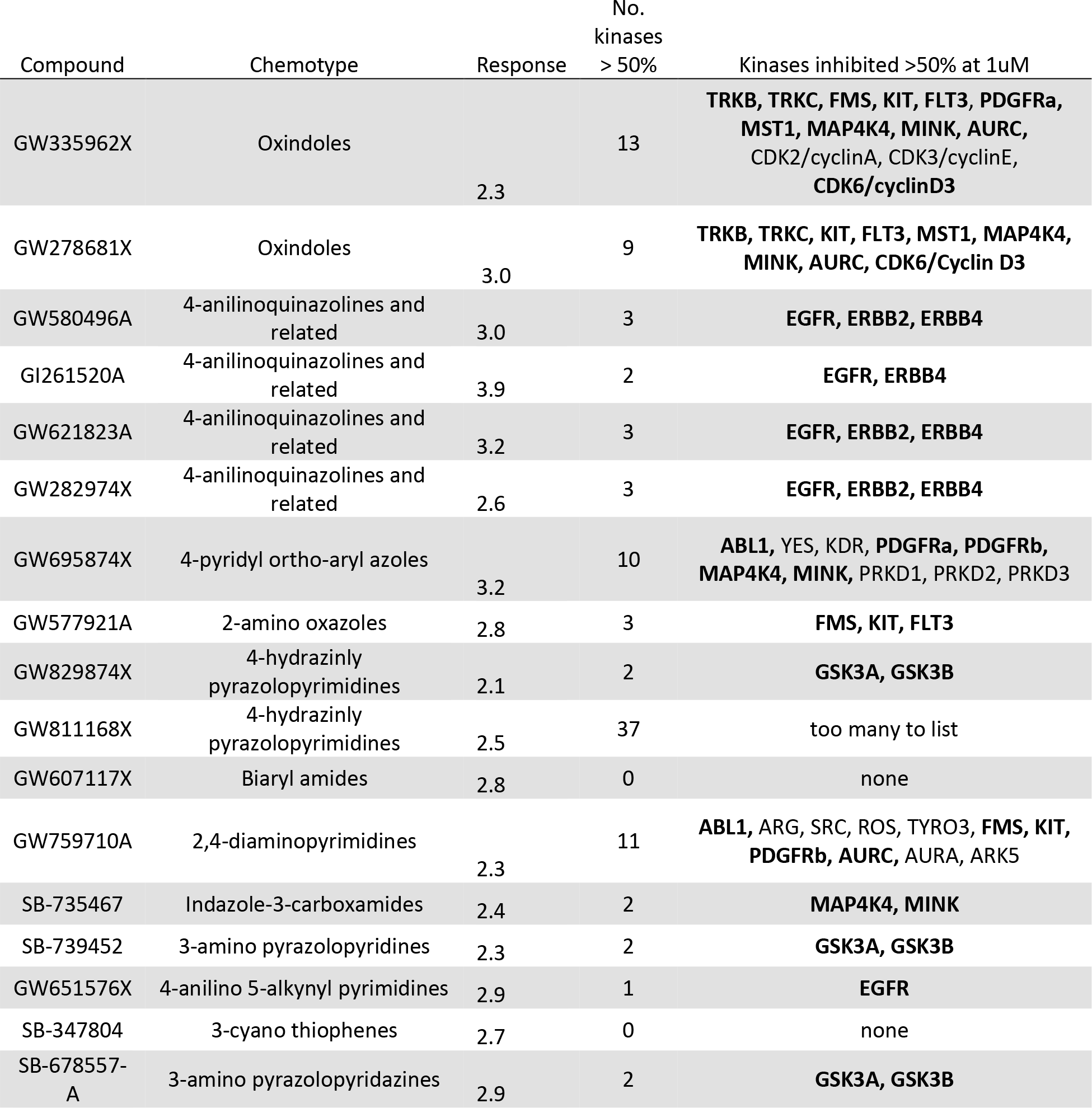
PKIS compounds that activated Nurr1 activity.

| Compound | Chemotype | Response | No.<br>kinases<br>> 50% | Kinases inhibited >50% at 1uM |
| --- | --- | --- | --- | --- |
| GW335962X | Oxindoles | 2.3 | 13 | **TRKB, TRKC, FMS, KIT, FLT3, PDGFRa, MST1, MAP4K4, MINK, AURC, CDK2/cyclinA, CDK3/cyclinE, CDK6/cyclinD3** |
| GW278681X | Oxindoles | 3.0 | 9 | **TRKB, TRKC, KIT, FLT3, MST1, MAP4K4, MINK, AURC, CDK6/Cyclin D3** |
| GW580496A | 4-anilinoquinazolines and related | 3.0 | 3 | **EGFR, ERBB2, ERBB4** |
| GI261520A | 4-anilinoquinazolines and related | 3.9 | 2 | **EGFR, ERBB4** |
| GW621823A | 4-anilinoquinazolines and related | 3.2 | 3 | **EGFR, ERBB2, ERBB4** |
| GW282974X | 4-anilinoquinazolines and related | 2.6 | 3 | **EGFR, ERBB2, ERBB4** |
| GW695874X | 4-pyridyl ortho-aryl azoles | 3.2 | 10 | **ABL1, YES, KDR, PDGFRa, PDGFRb, MAP4K4, MINK, PRKD1, PRKD2, PRKD3** |
| GW577921A | 2-amino oxazoles | 2.8 | 3 | **FMS, KIT, FLT3** |
| GW829874X | 4-hydrazinly pyrazolopyrimidines | 2.1 | 2 | **GSK3A, GSK3B** |
| GW811168X | 4-hydrazinly pyrazolopyrimidines | 2.5 | 37 | too many to list |
| GW607117X | Biaryl amides | 2.8 | 0 | none |
| GW759710A | 2,4-diaminopyrimidines | 2.3 | 11 | **ABL1, ARG, SRC, ROS, TYRO3, FMS, KIT, PDGFRb, AURC, AURA, ARK5** |
| SB-735467 | Indazole-3-carboxamides | 2.4 | 2 | **MAP4K4, MINK** |
| SB-739452 | 3-amino pyrazolopyridines | 2.3 | 2 | **GSK3A, GSK3B** |
| GW651576X | 4-anilino 5-alkynyl pyrimidines | 2.9 | 1 | **EGFR** |
| SB-347804 | 3-cyano thiophenes | 2.7 | 0 | none |
| SB-678557-A | 3-amino pyrazolopyridazines | 2.9 | 2 | **GSK3A, GSK3B** |

### Nurr1 Inhibitors

7 compounds inhibited Nurr1 activation without affecting the cellular ATP decrease caused by 6-MP (Figure 2 **and** Table 2). Of these seven compounds, there were three compounds represented with one chemotype (Benzimidazole_N-thiophenes). Some of these compounds had some overlapping kinase targets (highlighted in Table 2).

**Figure 2.**
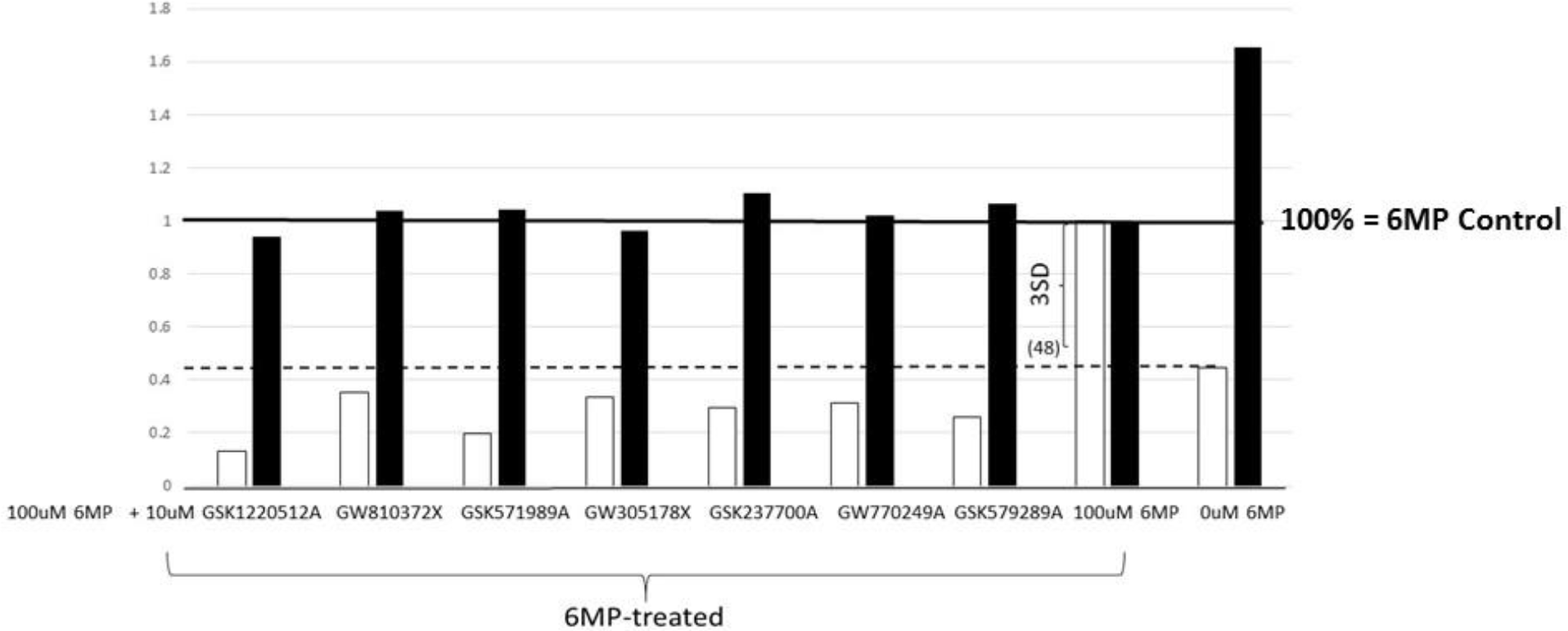
Seven compounds from PKIS screen inhibited 6MP-induced Nurr1 actiVity without. Significantly affecting the cellular ATP decrease caused by 6-MP. Controls are 100uM 6MP and OuM 6MP treatments. Data are normalized to the 100uM 6MP treatment (1.0 = 100%). 3 Standard deviations from the 6MP control valueis illustrated by bracket. White squares represent Nurr1 activity and Black squares represent cellular ATP levels.

**Table 2.** PKIS compounds that inhibited 6MP-induced Nurr1 activity and had no effect on cellular ATP levels.

| Compound | Chemotype | No. kinases<br>> 50% Inh | Kinases inhibited >50% at<br>1uM |
| --- | --- | --- | --- |
| **GSK1220512A** | 2,4-dianilino_pyrrolopyrimidines | 47 | too many to show |
| **GW305178X** | Oxindoles | 74 | too many to show |
| **GW810372X** | 2H-<br>3_pyrimidinyl_pyrazolopyridazines | 12 | PKY2, AurC, CDK2/CycE,<br>CDK2/CycA, GSK3A, GSK3B,<br>CLK2, HIPK1, HIPK4,<br>DYRK1B, DYRK1A |
| **GSK571989A** | Benzimidazole_N-thiophenes | 14 | MUSK, **PDGFRa, PDGFRb**,<br>MST1, **LOK, PLK1, ARK5**,<br>**BRSK1, BRSK2, PIM1, PIM3**,<br>NEK1, NEK2, **NEK9, PI3Kd** |
| **GSK237700A** | Benzimidazole_N-thiophenes | 11 | BLK, RET, KDR, FLT4,<br>**PDGFRa, PDGFRb, TNK1**,<br>LRRK2, **LOK, PLK1, BRSK2** |
| **GW770249A** | Fuopyrimidines_and_related | 50 | too many to show |
| **GSK579289A** | Benzimidazole_N-thiophenes | 10 | **PDGFRa, PDGFRb, LOK**,<br>**PLK1, PIM1, NEK9, ARK5**,<br>**BRSK1, BRSK2, PI3Kd** |

### Nurr1 Inhibitors

9 compounds inhibited Nurr1 and additionally inhibited the cellular ATP decrease caused by 6-MP (Figure 3 **and** Table 3). Assay of kinase activity data on the compounds showed that most of these compounds affected multiple kinase targets with most inhibiting more than 20 kinases above 50% Inhibition. Two of the hit compounds, GW284372X and GSK317315A, in contrast, only inhibited 2 and 3 kinases respectively above 50% Inhibition. These targets were not overlapping and further investigation would be required to identify the target(s) of this phenotype.

**Figure 3.**
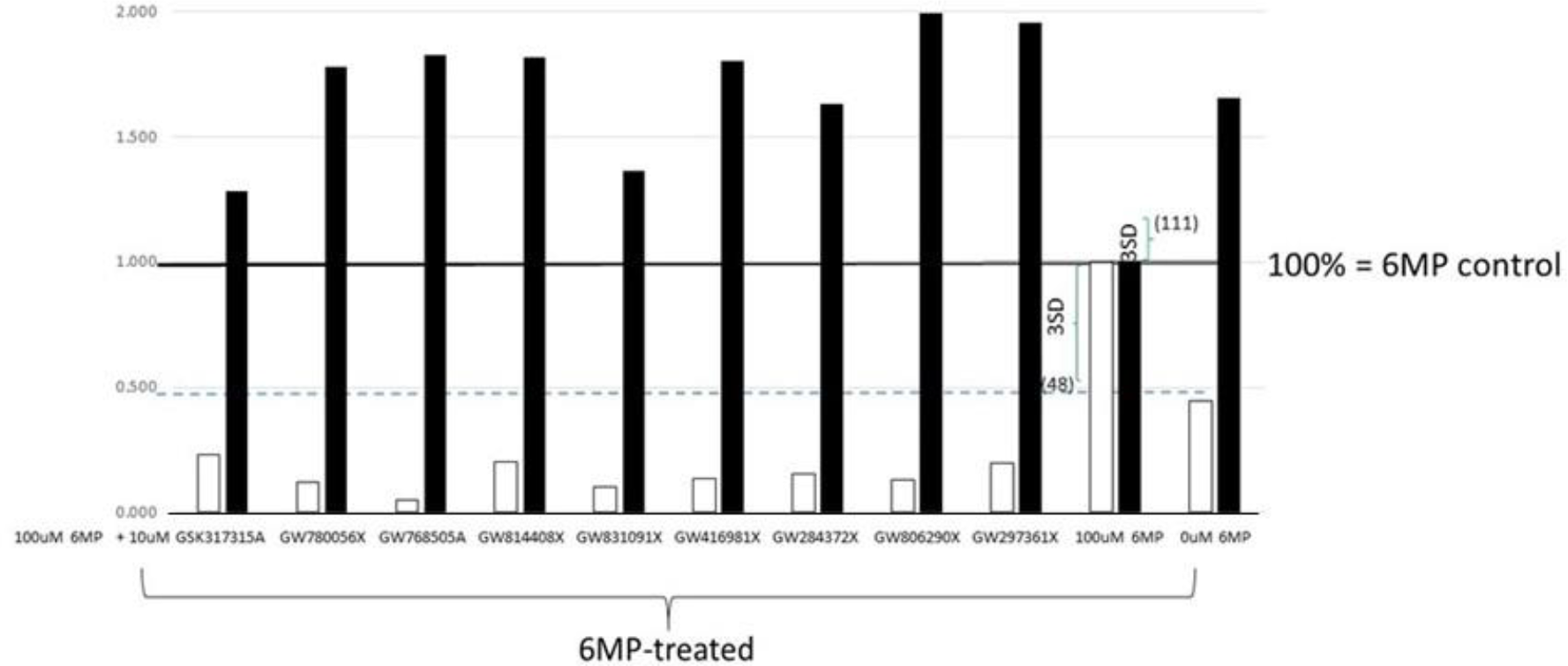
Nine compounds inhibited 6MP-induced Nurr1 activity and additionally inhibited the cellular ATP decrease caused by 6-MP. Controls are 100uM 6MP and OuM 6MP treatments. Data are normalized to the 100uM 6MP treatment (1.0 = 100%). 3 Standard deviations from the 6MP control value is illustrated by bracket. White squares represent Nurr1 activity and Blacks quares represent cellular ATP levels.

**Table 3.** PKIS compounds that inhibited 6MP-induced Nurr1 activity but had a significant effect on cellular ATP levels.

| Compound | Chemotype | No. kinases > 50% Inh | Response |
| --- | --- | --- | --- |
| GW806290X | 2H-3-pyrimidinyl pyrazolopyridazines | 41 | 1.99 |
| GW297361X | Oxindoles | 59 | 1.96 |
| GW768505A | Fuopyrimidines and related | 62 | 1.83 |
| GW814408X | 4-hydrazinly pyrazolopyrimidines | 45 | 1.81 |
| GW416981X | Oxindoles | 50 | 1.8 |
| GW780056X | 2H-3-pyrimidinyl pyrazolopyridazines | 34 | 1.75 |
| GW284372X | 4-anilinoquinazolines and related | 2 | 1.63 |
| GW831091X | 3-aminopyrazoles | 21 | 1.36 |
| GSK317315A | Benzimidazole N-thiophenes | 3 | 1.25 |

## Discussion

Nurr1 is a potentially important target for multiple diseases. We have carried out a primary screen of a broad set of kinase inhibitors using a cell-based assay developed using the amino terminal portion of the receptor. The screen identified multiple kinase inhibitors that increase or decrease activity of the transcriptional activity of Nurr1 in our assay. Assessment of potency of each of the identified hits is awaiting re-testing the compounds in dose-response assays. The dose-response assays will be conducted in the presence and absence of 6-MP to identify those molecules which are 6-MP independent. Additional assessment of the hits will also be carried out using full-length receptors to assure that regulation by the molecules is not restricted to the specific amino terminus construct used in our primary screen.

Our hypothesis is that these compounds are altering the phosphorylation status of the AF1 region of NR4A2, a key regulatory region of the receptor, altering the activity of an upstream kinase in the NR4A2 regulatory cascade. To support this hypothesis, additional experiments would be needed to test whether these compounds induce quantitative or qualitative changes in phosphorylation status of Nurr1 in cells.

Beyond their value as tool compounds in regulating Nurr1, the hits from our screen could potentially also aid in the identification of target kinases in the Nurr1 regulatory cascade. A broader “big data” analysis of Nurr1 activity data (both active and inactive compounds) cross-matched to their kinase selectivity will be carried out. Initial analysis of the data revealed that most of the kinase inhibitors identified have fairly broad selectivity profiles making it difficult to pinpoint specific candidate kinase targets at this time. Testing of analogues of the hits and also testing more recent versions of kinase inhibitor sets available from the SGC (https://pharmacy.unc.edu/research/sgc-unc/) could aid in furthering this analysis.

It is notable that two distinct NR4A2 inhibitor profiles were observed: compounds that inhibited 6-MP induced Nurr1 activity without altering cellular ATP levels and compounds that blocked the changes in 6-MP induced cellular ATP levels. It is possible that activation of the two sets of compounds activate NR4A2 by distinct mechanisms. It has been proposed previously that NR4A2 may be activated by generalized cellular stress such as decreased purine levels (Ordentlich *et al*., 2003).

In sum, these compounds serve as starting points to dissect the NR4A2 signaling pathway and identify tractable targets within the NR4A2 activation pathway that could be exploited for drug discovery. To identify the best tool compounds for pathway experiments, further characterization of these compounds is needed.

